# Challenging a preconception: Optoacoustic spectrum differs from the absorption spectrum of proteins and dyes for molecular imaging

**DOI:** 10.1101/2020.02.01.930230

**Authors:** Juan Pablo Fuenzalida Werner, Yuanhui Huang, Kanuj Mishra, Robert Janowski, Paul Vetschera, Andriy Chmyrov, Dierk Niessing, Vasilis Ntziachristos, Andre C. Stiel

## Abstract

Optoacoustic (photoacoustic) imaging has seen marked technological advances in detection and data analysis, but there is less progress in understanding the photophysics of optoacoustic signal generation of commonly used contrast agents, such as dyes and chromoproteins. This gap blocks the precise development of novel agents and the accurate analysis and interpretation of Multispectral Optoacoustic Tomography (MSOT) images. To close it, we developed a multimodal laser spectrometer (MLS) to enable the simultaneous measurement of optoacoustic, absorbance, and fluorescence spectra. MLS provides reproducible, high-quality optoacoustic (non-radiative) spectra by using correction and referencing workflow. Herein, we employ MLS to analyze several common dyes (Methylene Blue, Rhodamine 800, Alexa Fluor 750, IRDye 800CW and Indocyanine green) and proteins (sfGFP, mCherry, mKate, HcRed, iRFP720 and smURFP) and shed light on their internal conversion properties. Our data shows that the optical absorption spectra do not correlate with the optoacoustic spectra for the majority of the analytes. We determine that for dyes, the transition underlying the high energy shoulder, which mostly correlates with an aggregation state of the dyes, has significantly more optoacoustic signal generation efficiency than the monomer transition. Our analyses for proteins point to a favored vibrational relaxation and optoacoustic signal generation that stems from the neutral or zwitterionic chromophores. We were able to crystalize HcRed in its optoacoustic state, confirming the change isomerization respect to its fluorescence state. Such data is highly relevant for the engineering of tailored contrast agents for optoacoustic imaging. Furthermore, discrepancies between absorption and optoacoustic spectra underline the importance of correct spectral information as a prerequisite for the spectral-unmixing schemes that are often required for *in vivo* imaging. Finally, optoacoustic spectra of some of the most commonly used proteins and dyes in optical imaging, recorded on our MLS, reveal previously unknown photophysical characteristics, such as unobserved photo-switching behavior.

## INTRODUCTION

Optoacoustic (OA, also termed photoacoustic) imaging visualizes optical contrast with ultrasound resolution, enabling resolved, real-time *in vivo* imaging well-beyond the 1 mm penetration depth of typical optical methods (Ntziachristos, 2010). Therefore, it is emerging as a particularly appealing *in vivo* imaging method to study tumor biology (Omar et al., 2015), inflammatory diseases (Aguirre et al., 2017), developmental processes (Vetschera et al., 2019), or brain functioning in mammals *in vivo* (Ovsepian et al., 2017). However, in the majority of such studies, OA imaging relies entirely on endogenous absorbing-agents, like blood-hemoglobin, melanin, or lipids. In contrast, targeted labels enable longitudinal, cell-specific *in vivo* imaging, drastically expanding the potential applications of OA imaging (Razansky et al., 2009). Several potential agents have been tested for OA imaging, including nanoparticles and targeted dyes (reviewed in Gujrati *et al*., 2017(Gujrati et al., 2017)), as well as transgenic labels, such as fluorescent proteins, bacteriophytochromes, and their switchable variants (reviewed in Brunker *et al*., 2017(Brunker et al., 2017)).

In some cases, these labels were specifically developed for OA applications (Chee et al., 2018; Li et al., 2016). However, despite these successes, most label choices and the strategies for their development are still based on OA performance and photophysics inferred from independent transmission-mode absorption and fluorescence spectroscopy data. As shown by our previous work and others, low quantum yield and high molar absorption coefficient do not automatically translate to high optoacoustic signal(Fuenzalida Werner et al., 2019).

In this work, we sought to develop a method of spectral analysis that combines absorption and fluorescence with OA spectral data to enable the full characterization of photophysical parameters of OA labels. Such a method would help to identify competing transitions (e.g., triplet states, long-lived dark states, or molecular isomerization; see **Figure 1a**). This data is a key to accelerate the tailored development of OA contrast agents. Furthermore, setting standards and building a reliable database of spectral information could guide researchers in their choice of suitable labels, similar to the “molecular brightness” standard used in fluorescence imaging. Lastly, a combined method of spectral analysis would enable the study of various photophysical phenomena, for example, dye aggregation, exciton coupling, and energy conversion.

**Figure 1.**
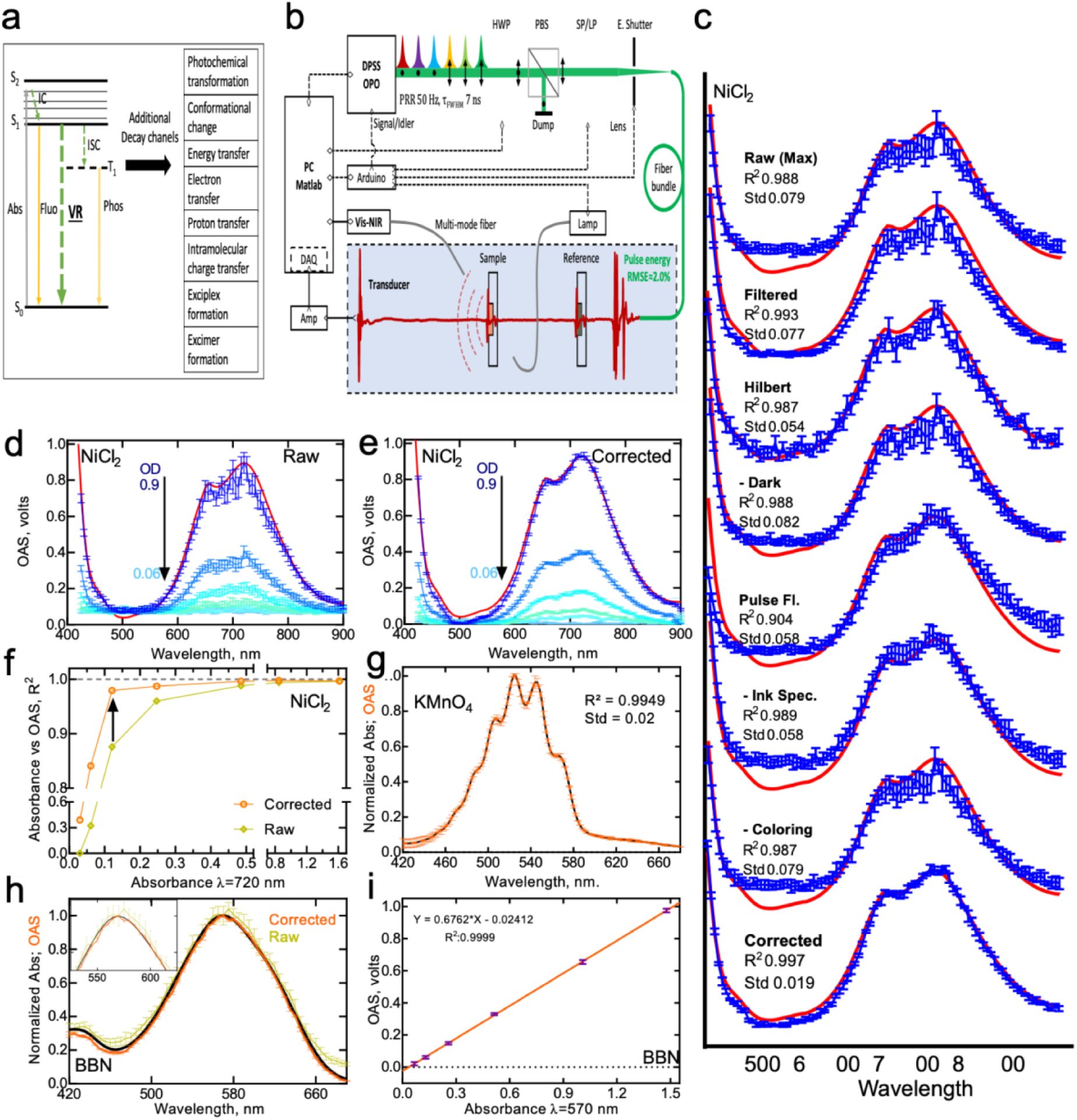
**(a)** Photophysics of OA signal generation. Jablonski energy diagram. Plain lines represent radiative transitions, dashed lines represent nonradiative processes. Abs: absorbance S0: ground state, S1: singlet excited state, T1: primary triplet excited state, VR: vibrational relaxation ISC: intersystem crossing, IC: internal conversion, Pho: phosphorescence, Fluo Fluorescence. **(b)** Setup diagram (explanations and abbreviations can be found in the text). **(c)** Improvement in optoacoustic spectrum of NiCl_2_ brought by each of the correction steps. The coefficient of determination R_2_ and standard deviation (Std) between measurements are shown to indicate the improvement by each of steps. ‘Max (Raw)’ shows a spectrum that can be obtained using simply the maximum values in sample optoacoustic signals generated by homogenized laser energy across wavelength. ‘Filtered’ shows the spectrum can be obtained using filtering techniques on signals. ‘Hilbert’ shows the spectrum formed by maximum values of the signal envelops using Hilbert transform. ‘-Dark’ shows the spectrum that is without DC bias by subtracting values measured in dark (no laser). ‘Pulse Fl.’ shows the spectrum after pulse-to-pulse fluctuation correction enabled by our in-line reference. ‘-Ink Spec.’ shows the spectrum after removing the ink spectrum which is induced when performing fluctuation correction. ‘-Coloring’ shows the spectrum after the correction for the light fluence coloring effect induced by the reference Ink and US-couplant water. ‘Corrected’ shows the spectrum that can be obtained after performing all the afore-mentioned steps (further explanations can be found in the text). **(d)** Raw optoacoustic spectrum of NiCl_2_ at different concentrations and **(e)** after correction. **(f)** The coefficients of determination of linearity between absorbance and optoacoustic spectrum of NiCl_2_ at decreasing concentrations. **(g)** OA and absorbance spectrum for potassium permanganate at 2 nm steps. **(h)**Raw and corrected optoacoustic spectrum of BBN. **(i)** Linear relation between absorbance and OA signal for BBN.

We introduce herein a multi-modal laser spectrometer (MLS) that allows us to measure optoacoustic spectra with high precision and spectral resolution, concomitantly with absorption, and fluorescence spectra. Key characteristics of this system are i) illumination with pulsed lasers, similar to OA imaging; ii) homogenization of fluence for all measured wavelengths; iii) in-line reference and correction to overcome laser instabilities; iv) simultaneous detection of fluorescence excited by the laser pulse, and v) simultaneous absorbance measurement. In contrast to other devices for recording OA spectra, which required samples with high concentrations and/or large volumes (Laufer et al., 2013; Schneider and Coufal, 1982; Teng and Royce, 1980), our MLS requires only 200 μL of minimum 0.2 optical density (OD) concentration for high-quality, reproducible spectra (R_2_ > 0.95). This high spectral quality allowed us the study of a range of dyes namely, Methylene Blue, Rhodamine 800, Alexa Fluor 750, IRDye 800CW and Indocyanine green and chromoproteins namely, sfGFP, mCherry, mKate, HcRed, iRFP720 and smURFP. In most cases, we show significant differences between absorbance and OA spectra. We correlated these discrepancies either to an aggregation state of the dyes or to the isomeric state of the chromophores. Finally, we discuss how such high-quality MLS spectra can empower future optoacoustic contrast agent development and spectral unmixing to achieve high detection sensitivity in molecular imaging.

## RESULTS & DISCUSSION

### Multi-modality laser spectrometer (MLS)

In order to fully characterize the photophysical parameters underlying the OA signal of a particular sample, it is essential to obtain exact measurements of absorbance, fluorescence, and OA spectra under the same light fluency to assess the relative contributions of the different electronic transitions. Based on expertise gained measuring the OA kinetic parameters of photo-controllable proteins (Vetschera et al., 2018), we developed a multi-modal laser spectrometer (MLS, **Figure 1b**) that can record all three modalities simultaneously in the wavelength range between 420-1000 nm. In brief (**Figure 1b**, see Material and Methods details), for the OA and fluorescence spectroscopic measurements, we used an optical parametric oscillator (OPO) laser with a 50 Hz repetition rate and average 7 ns pulse width as the light source. The laser energy at different wavelengths (420-1000 nm) was kept constant by controlling a motor-controlled half-wave plate (HWP) in combination with a polarizing beam-splitter (PBS) according to a lookup table of pulse energies at all wavelengths. The laser energy was measured by an in-line power meter before every measurement. The light was delivered to the sample submerged in water through a fiber bundle. The light travels first through a 200 μm thick in-line reference chamber filled with Indian ink (0.2-0.7 OD) before reaching a similar chamber containing the actual sample. The absorbance of ink was determined relative to the absorbance of the sample in order to optimize the dynamic range of our data acquisition system. The laser excited optoacoustic wave is recorded using a cylindrically focused ultrasound transducer with a central frequency of 3.5 MHz. The signal is firstly amplified by a variable-gain amplifier (Amp) and then digitized by a 12-bit digitizer (DAQ) and PC. The fluorescence signal, which is excited by the same laser pulse, is recorded simultaneously by a fiber-coupled diode-array spectrometer oriented 45° to the laser beam path. The absorption is recorded using the same spectrometer an instant after the OA and fluorescence measurements by blocking the laser beam with an electronically controlled shutter (E. Shutter) and opening the shutter of a halogen lamp to illuminate the sample with a fiber opposite to the spectrometer.

An essential part of the MLS performance is a stepwise correction procedure enabling highly accurate OA spectral measurements. In brief (**Figure 1c**, Supplementary **Figure 1–3**, see Material and Methods details): An initial laser energy look-up table is computed to correct for the major wavelength-dependent changes in laser energy. This table is then used to control the HWP and beam splitter to maintain the same energy at each wavelength. We adopted a measure- and-fit routine to measure the lookup table and designed an iteration-based lookup table update method. This afforded a fast lookup table measurement and precise control of the laser energy. However, the response of the power meter is too low to ensure the correction of pulse-to-pulse energy fluctuations. Furthermore, while the peak power variations might be correctible with a fast-response power meter, the variation in pulse width is harder to measure and correct with high-precision. Thus, for pulse-to-pulse energy fluctuation correction, we rely on the in-line Indian ink-reference to deliver a time-shifted signal excited by the same pulse as the actual sample. This method of correction necessitates the subtraction of the Indian ink absorbance spectrum from the fluctuation corrected spectrum. We further corrected for the spectral coloring induced by the in-line reference material (Indian ink) and the water path of laser, to reduce possible fluence alterations.

### Characterization of the MLS

To validate the optics setup and the correction steps, we analyzed the spectra of NiCl_2_. NiCl_2_ is known to have no apparent photo-fatigue and a complete transfer of absorbed energy into acoustic waves via non-radiative decay channels, so its absorption spectrum should correlate with its optoacoustic spectrum (Laufer et al., 2010). The OA spectra of NiCl_2_ measured without any correction (R_2_ 0.988 and standard deviation (SD) 7.9%) and using the full correction protocol of our MLS (R_2_ 0.997 and SD 1.9%) demonstrate the increase of spectral quality, with each correction step (**Figure 1c**). The improvement of spectral quality through our measurement protocol is significant over a range of concentrations, with an R_2_ of 0.85 for a NiCl_2_ sample with OD as low as 0.05 (**Figure 1d to f**). In order to test the wavelength accuracy of our setup, we measured OA and absorbance spectra of potassium permanganate (**Figure 1g**), and we were able to match its six absorption peaks with 2 nm accuracy correctly. To further characterize our MLS system, we similarly measured different concentrations of Brilliant Black N dissolved in PBS (BBN, **Figure 1h, and i**). BBN, similar to NiCl_2_, has a negligible fluorescence quantum yield (QYfluo) and very high photostability, and thus undergoes fully non-radiative deexcitation. Importantly, BBN, in contrast to NiCl_2_, is soluble and stable in aqueous buffer solutions. BBN thus shares a common Grüneisen parameter (Abbruzzetti et al., 1999; Brown et al., 2015; Laufer et al., 2010) with labels for *in vivo* imaging, which are frequently dissolved in aqueous buffer systems. The combination of the excellent solubility of BBN and its stable generation of OA signals over a range of concentrations and excitation energies makes it well suited to serve as a universal standard for precise and reproducible OA measurements. Such allows reproducible characterization of dyes and other labels eventually allowing the determination and reporting of universally meaningful OA signal strength values.

### OA signal generation efficiency

Vibrational relaxation (non-radiative energy decay) from an excited state produces a localized change in temperature, resulting in an optoacoustic signal (OAS, **Figure 1a**) via thermal expansion. This change in pressure is generally described by equation (1) (Deán-Ben et al., 2017).

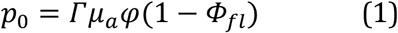

In equation 1, *μ_a_* is the local optical absorption coefficient, *φ* is the local light fluence, and *Φ_fl_* is the fluorescence quantum yield. *Γ* is the dimensionless Grüneisen parameter, which represents the thermoelastic properties of the medium. Equation 1 dismisses all other competitive channels, like conformational changes, photochemical transformations, intramolecular charge transfer, electron transfer, intersystem crossing, proton transfer, exciplex formation, excimer formation, and energy transfer (Land, 1979) (**Figure 1a**). All of these processes reduce the number of available electrons that undergo vibrational relaxation and produce a change in pressure in the time (Mishra et al., 2019; S. P. McGlynn, 1969; Schaberle et al., 2016). Thus, in order to accurately compare different optoacoustic capacities, the OAS has to be normalized by the absorbance, so that the slope of the linear relation between the OAS and absorbance yields the photoacoustic generation efficiency (PGE), which is a measure of how much of the absorbed energy is converted to pressure (Abbruzzetti et al., 1999; Braslavsky et al., 1983a; Brown et al., 2015; Laufer et al., 2013; Rosencwaig, 1980). As mentioned before, photostability and the lack of any fluorescence make BBN an ideal reference for photoacoustic measurements; accordingly, we assign it a PGE value of 1.0, assuming that no other decay channels take place (Abbruzzetti et al., 1999; Brown et al., 2015). After BBN normalization equation one can be rewritten to derive equation 2. Thus, PGE is attenuated by fluorescence (*Φ_fl_*), and the energy ‘stored’ in long-lived transients such as a triplet state (*Φ_st_*) or any other photochemical decay pathway (*Φ_x_*)(Borges Dos Santos et al., 1999; Rios et al., 2007).

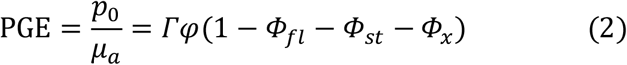

### Characterization of dyes

The sensitivity of some natural chromophores and organic dyes to aggregation and other environmental changes has been exploited for various sensing applications in OA imaging (Jiang and Pu, 2017; Lovell et al., 2011; Morgounova et al., 2013; Peters et al., 2019; Wang et al., 2018). However, the photophysics underlying the changes in the OA signals of these chromophores have never been thoroughly understood or described, precluding efficient and rational exploitation of such effects for the design of next-generation functional OA labels. Methylene Blue (MB) and Rhodamine 800 (Rh800) are two well-known synthetic xanthene dyes. MB has been commonly employed as a label in OA due to its lack of fluorescence (Taruttis and Ntziachristos, 2015). Rh800 would be even more favorable for OA imaging due to its more pronounced red absorption (682 nm vs. 660 nm); however, its PGE is attenuated by strong fluorescence at 712 nm (Alessi et al., 2013). Both dyes have the tendency to form aggregates through strong aromatic interactions. The aggregates have photophysical characteristics that are clearly distinguishable from those of the monomers based on additional spectral bands next to the monomers’ absorbance.

The H-aggregate, with the dye molecules stacked in the same orientation, shows a blue-shifted absorbance, while the J-aggregate, in which the molecules are shifted with respect to each other, shows a more red-shifted absorbance (Kasha et al., 1965). The formation of H-type aggregates commonly reduce fluorescence significantly, while J-type aggregates can increase fluorescence (Hestand and Spano, 2017, 2018; Kasha et al., 1965). Intersystem crossing (ISC) rates are also determined by the supramolecular architecture and orientation geometry of the aggregates (Hestand and Spano, 2017; Kasha et al., 1965). In general, the aggregation behavior is strongly depending on environmental conditions like pH, ionic strength, and crowding agents (Krieger et al., 2017; Moreno-Villoslada et al., 2010a). In order to create a commonly defined medium for our measurements that resemble the cellular environment, we dissolved the dyes in 10% fetal bovine serum (FBS) in phosphate buffer saline (PBS) (Bongsu Jung et al., 2014). In this environment, all dyes are in a mixed aggregation state, which allows us to study the different characteristics of the aggregate related transitions. Even more critically, this provides stable environments that enable us to observe the spectra without changes over time.

For solutions of MB with concentrations ranging from 17 to 29 μM, we found fluorescence emission only from excitation of the red-shifted spectral band, corresponding to the weakly fluorescent monomer, while the blue-shifted band, associated predominantly with the H-aggregates, did not show fluorescent transitions (**Figure 2a**). Conversely, the OA spectrum shows a comparably stronger signal for the H-aggregate band (610 nm), reflecting a higher OA signal generation (thus PGE) for this transition compared to the monomer (slope of 0.61 at 610 nm vs. 0.37 at 665 nm in **Figure 2b**). It is generally accepted that the loss of fluorescence in highly symmetric H-aggregates, like the one formed by MB, stems from a favored ISC (Kasha et al., 1965; Morgounova et al., 2013) (**Supplementary Figure 4**). This is corroborated by our photo-fatigue analysis of the different transitions, which shows faster bleaching for the H-aggregate at 610 nm (**Supplementary Figure 5**) than for the monomer at 665 nm. However, the vibronic relaxation following ISC, regularly occurring in the long microseconds range, was considered too slow to contribute to detectable OA signals(Visser et al., 2002) in our observation time window of sub-microseconds range, and thus would not explain the higher PGE of the H-aggregate. Interestingly, the ISC for MB is very fast (ns vs. μs) (Morgounova et al., 2013), suggesting that we can indeed detect the resultant vibronic relaxations. Consequently, we can show here that the formation of H-aggregates by MB results in fast ISC, which contributes to the observed strengthening of the OA signal.

**Figure 2.**
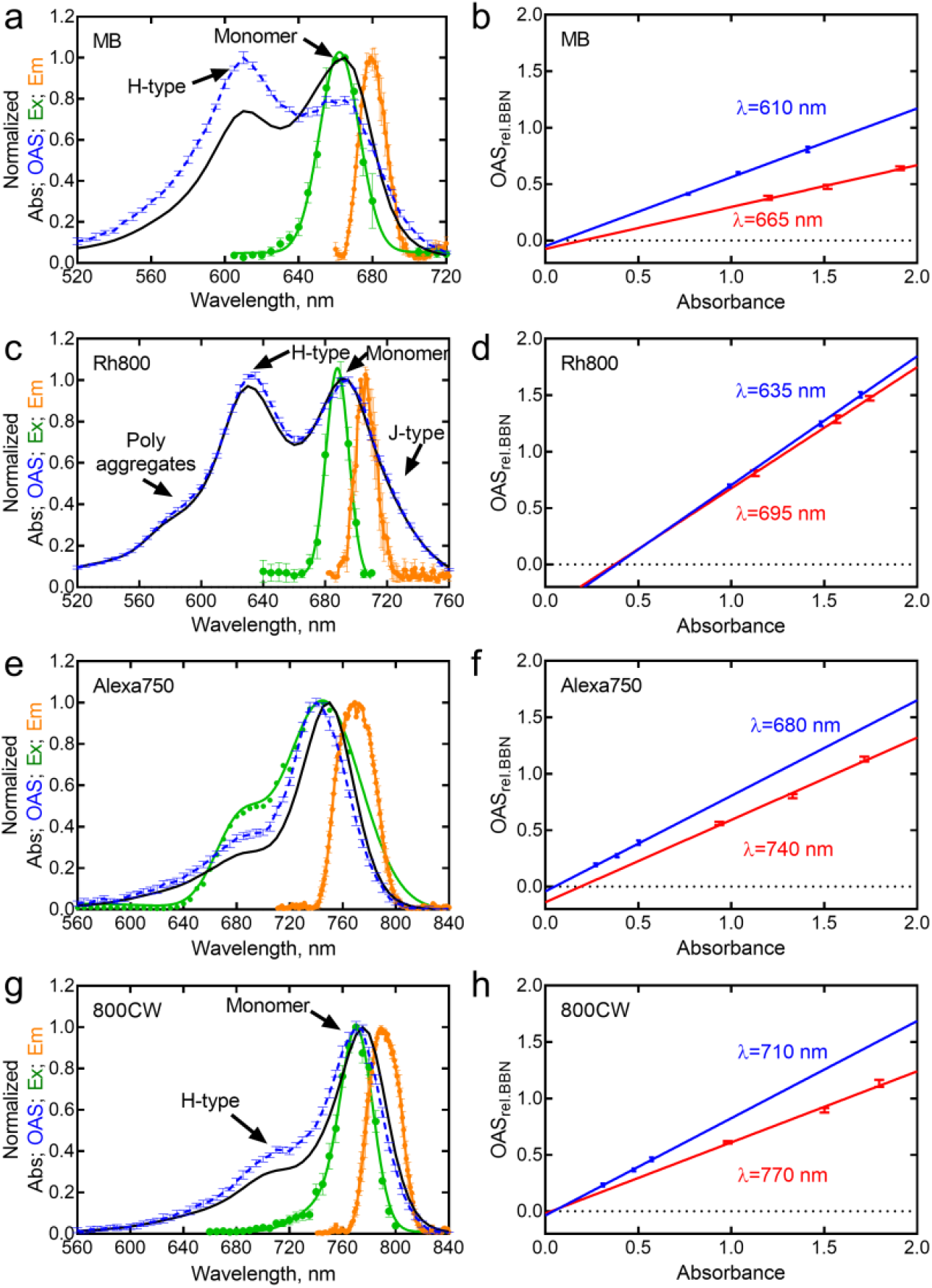
Normalized optoacoustic signal (OAS), absorption (Abs), emission (Em) and excitation (Ex) spectra for **(a)** MB - Methylene blue, **(c)** Rh800 - Rhodamine 800, **(e)** Alexa780 - Alexa Fluor 750 and **(g)** 800CW - IRDye 800CW. **(b,d,f,h)** The linear relationship of OAS and absorbance are given for the main peaks and shoulders, respectively.

Because the dye Rh800 fluoresces, it has thus far not been considered for applications in OA; however, the increase in the PGE of MB that we observed upon its aggregation prompted us also to study the photophysics of Rh800. We measured Rh800 in a concentration range from 30 to 49 μM. Rh800 shows a similar combination of monomer (695nm) and predominantly H-aggregate (635 nm) bands, similar to MB (**Figure 2c**). However, in contrast to MB, the PGEs of the two bands are only slightly different (1.14 and 1.08, **Figure 2d**). To our surprise, the PGE of the monomer form is high, despite its fluorescence, even surpassing our standard BBN (**Figure 1h and i**). Moreover, the bleaching rates of both transitions are moderate and suggest no strong ISC, as also observed for MB (**Supplementary Table 1, Supplementary Figure 5**). Upon closer inspection of the spectral relations, it becomes apparent that the fluorescence excitation spectrum is shifted in respect to the absorbance band at 695 nm, hinting at a third transition, presumably a non-fluorescent J-aggregate, in the flanks of the peak primarily attributed to monomer (720 nm, arrow). Further inspection of the spectra reveals an additional shoulder in the blue region (580 nm, **Figure 2c arrow**). Such high energy transitions in xanthene dyes are often correlated with higher-order aggregates and have been previously observed (Moreno-Villoslada et al., 2010b). This is additionally underpinned by a pronounced H-aggregate band suggesting that most of the dye is in an aggregate state. The acoustic wave for Rh800 at 695nm and 635nm differ from each other and they differ from the one of BBN (**Supplementary Figure 6**). It is likely that this change of the acoustic wave is directly related to the higher order aggregates. First, the aggregates represent a larger emitter changing the waveform and, second, the aggregates displace the water around the individual dye monomers and thus change the Grüneisen parameter. Both mechanisms favor the OA signals upon deexcitation, thus explaining the unusually high PGE. Moreover, the low bleaching rate (10% signal loss after 18 exaphotons (**Supplementary Figure 5**), suggests only minor contributions from transitions competing with the non-radiative decay (like fluorescence or ISC), further explaining the high PGE.

Alexa Fluor 750 (Alexa750) and IRDye 800CW (800CW) show absorbance peaks centered in the favorable blood window (749 nm and 774 nm, **Figure 2e, and g**), which is why they have been developed for *in vivo* fluorescence imaging applications. Despite their relatively high fluorescence (QY_fluo_ = 0.12 for both Alexa750 and 800CW), both dyes have already been used in OA imaging (Chen et al., 2018; Lakshman and Needles, 2015). Analysis of the spectra of Alexa750 shows a shift for the excitation maximum as well as the PGE maximum of 10 nm relative to the absorbance spectrum. This points to a transition at ca. 698 nm (**Figure 2e**), blue shoulder, arrow A). Additionally, this shoulder bleaches considerably less than the main peak, with only 40% loss versus 80% after 60000 light pulses for the shoulder and main peak, respectively (**Supplementary Figure 5**). This low bleaching and effective deexcitation in comparison to the main peak of Alexa750 points to a strong triplet tendency of the main peak leaving the transition in the shoulder as the more effective one for imaging. Similar to the xanthene dyes, the absorbance spectrum of 800CW has two peaks (**Figure 2g**) with one centered at 710 nm and the second one centered at 770 nm. The fluorescence excitation spectrum mirrors the primary absorption peak (770 nm), with a stronger PGE in the blue shoulder (710 nm), attributable to H-aggregates. Despite evidence of 800CW showing excitation coupling with other aromatic dyes (Joseph et al., 2017; Zhang et al., 2017), there is no description in the literature of aggregation of 800CW molecules. The observed behavior, spectrally similar to MB and Rh800 described above, might point to the blue shoulder resulting from H-type aggregation for 800CW. Intuitively, the formation of self-aggregates in 800CW seems to be unfavorable due to 800CW’s strong negative character (four negatively charged sulfonate groups). However, it has been noted that several other similarly charged dyes undergo aggregation and exciton coupling in the presence of macromolecules, such as polymers, HSA, or BSA (Hoffmann et al., 2016; Liang-Schenkelberg et al., 2017). Thus, the FBS that is present in our solutions could lead to the formation of H-aggregates of 800CW, explaining the observed exciton coupling behavior.

### Indocyanine green

Indocyanine green (ICG) is a dye widely used in OA due to its relatively strong signal intensity, favorable absorbance at 800 nm, and FDA approval (Beziere et al., 2015; Kim et al., 2010; Weber et al., 2016; Wilson et al., 2018). The zwitterionic and hydrophobic character of ICG allows its aggregation in an aqueous solvent at very low concentrations (Philip et al., 1996; Zweck and Penzkofer, 2001). Furthermore, it has been shown that proteins, such as albumin, can modify the aggregation state of ICG (Bongsu Jung et al., 2014; Philip et al., 1996). Similar to the previous dyes, we prepared and measured solutions of ICG with concentrations ranging from 2 to 26 μM, in 10% FBS. OA and absorbance spectra show a constant mismatch (**Figure 3 a and b**, arrow), with the whole spectral shape exhibiting a redshift and an increase of the 745 nm band with increasing concentration. The PGE for the OA shoulder centered at 745 is 0.80. Interestingly the absorbance at 795 nm does not correlate linearly with the OAS. To compensate for this non-linear behavior, we used two windows for the calculation of the PGEs, allowing a linear fit (**Figure 3c**). The PGE at 795 nm (0.62) is significantly lower at concentrations below 10 μM than it is above 10 μM (0.87), suggesting a higher PGE for aggregates of ICG. Thus, the difference between the dimer/monomer ratio from absorbance and OA represents the aggregation tendency of ICG (**Figure 3d**). The inflection point of the sigmoidal curve at 9.2 μM is between the two linear regimes chosen for PGE calculation. The PGE increases at the monomer peak (795 nm) with increasing concentration because the fraction of ICG aggregates also absorb at this wavelength. The discrepancies between absorbance and OA spectra observed could arise from different species of ICG present primarily at low concentrations, such as monomer, monomer bound to proteins (FBS), dimer, and dimer bound to proteins, as previously suggested (Philip et al., 1996). At higher concentrations, aggregated ICG interacting with proteins becomes the predominant species, the acoustic and absorption spectra of which do not vary significantly. Our results for ICG prove clearly that aggregation increases optoacoustic signal generation. Beyond encouraging the production of nanoparticles with highly aggregated ICG for optoacoustic imaging, this also suggests the use of the aggregation states as ratiometric readout.

**Figure 3.**
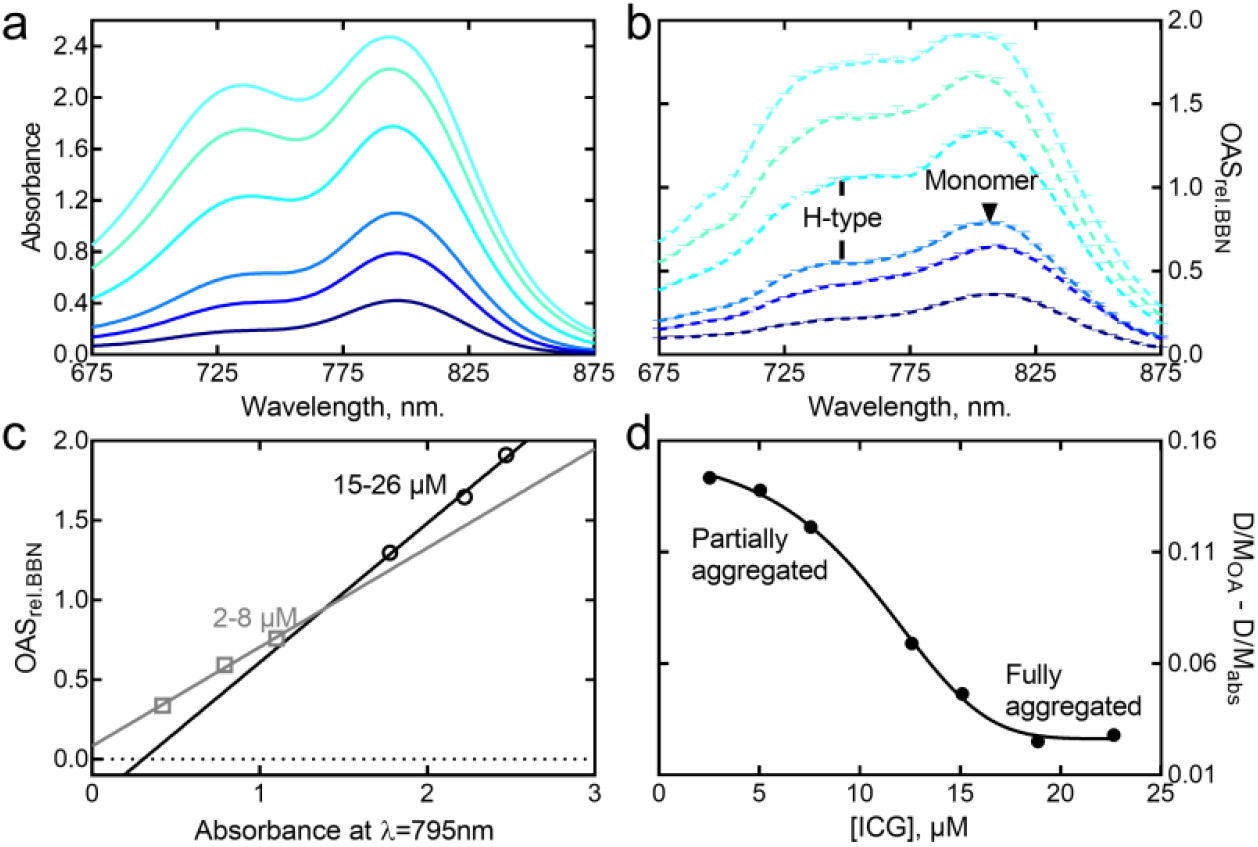
**(a and b)** Absorption and optoacoustic spectrum of ICG at increasing concentration. **(c)** Segmented linear relation between absorbance and optoacoustic signal for ICG. **(d)** Difference between the dimer/monomer ratio from absorbance and OA at increasing ICG concentration.

### Characterization of GFP-type chromophore bearing proteins

Genetically encode labels are a prerequisite for longitudinal imaging and one of the reasons for fluorescence imaging becoming a standard tool in life sciences. GFP-like labels are prominent in fluorescence imaging and, despite their mostly visible absorbance profiles, have also been studied and employed as OA labels (Filonov et al., 2012; Laufer et al., 2013; Liu et al., 2016; Razansky et al., 2009; Scheibe et al., 2019). However, biliverdin bearing chromoproteins like Bacteriophytochromes (BphPs) or Phycobiliproteins (Pbp) are advantageous due to their near-infrared (NIR) absorbance and have already been employed prominently in OA imaging (Chee et al., 2018; Filonov et al., 2012; Fuenzalida Werner et al., 2019; Li et al., 2018; Märk et al., 2018; Yao et al., 2015). The hydroxybenzylidene imidazolidone chromophores of GFP-like proteins share complex photophysics involving different protonation states of the chromophore (neutral, zwitterionic, and anionic) and a plethora of additional effects like excited-state proton transfer (ESPT) and long-lived dark states (often involving isomerization along the methine bridge). For a review, see Van Thor *et al*., 2009(van Thor, 2009).

We focused our spectral analysis on sfGFP (an advanced variant of the original *Aequorea victoria* GFP (Pédelacq et al., 2006)), red mKate2 (Shcherbo et al., 2009), mCherry (Shaner et al., 2004), tdTomato (Shaner et al., 2004), and far-red non-fluorescent HcRed (Gurskaya et al., 2001). Comparing the OA and absorption spectra of all these proteins reveals that in almost all cases (except for tdTomato), the blue shoulder of the peak is pronounced in OA, sometimes even leading to a blue-shift of the OA maxima compared to the absorption. Specifically, sfGFP shows a maximum for absorbance and OA at 480 nm; however, the shoulder in the blue region of the OA spectra is more prominent, including an identifiable slope rising at ~425 nm for the neutral chromophore (**Figure 4a and b**). This is also reflected in a higher PGE for the blue transition compared to the main peak (425 nm: 0.75; 480 nm: 0.34). mKate2 has an optoacoustic spectrum that is 5 nm blue-shifted for the absorption spectra (585 nm vs. 580 nm) and matching PGEs of 550 nm: 0.64 and 580 nm: 0.51 (**Figure 4c and d**). For mCherry, the maximum is similarly blue-shifted compared to the absorbance (580 nm vs. 585 nm), but with the corresponding PGE only slightly higher for the blue-shifted peak than for the absorbance peak (545 nm: 0.54; 585 nm: 0.53, **Figure 4e and f**). For the far-red and non-fluorescent tetrameric protein HcRed, the optoacoustic spectrum is blue-shifted by 20 nm with respect to the absorption spectrum, with the peak center at 570 nm instead of 590 nm (**Figure 4g and h**). Accordingly, the PGEs at 570 nm and 590 nm are 0.97 and 0.80, respectively. In general, except again for tdTomato, the PGEs of the main peaks are in good agreement with reported fluorescent quantum yields (**Supplementary Table 1**). tdTomato, on the other hand, a protein related to mCherry, is the only example of a protein measured in our study that shows an OA spectrum that is highly displaced to the red - a maximum in the OA spectrum at 565 nm, rather than 555 nm, with a PGE of 0.79 at 485 nm and of 0.83 at 565 nm (**Figure 4i and j**).

**Figure 4.**
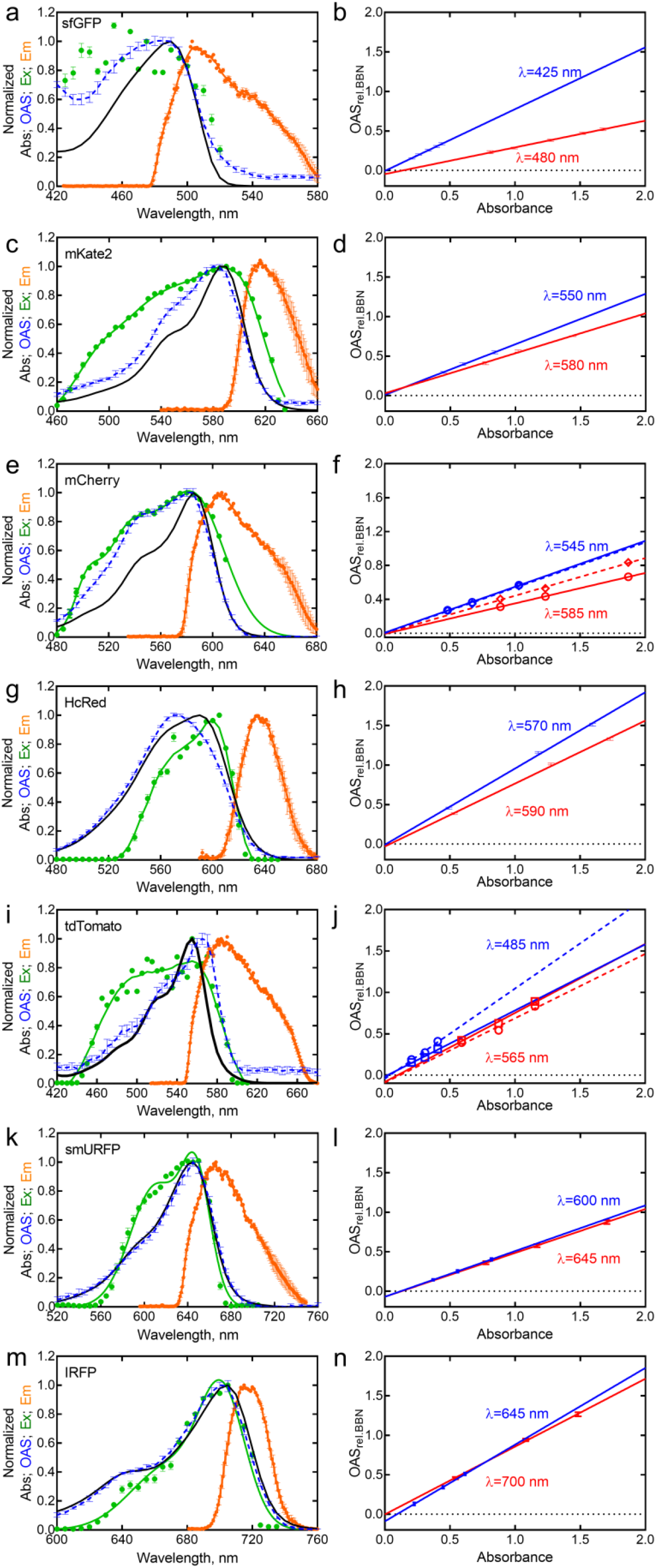
Normalized optoacoustic signal (OAS), absorption (Abs), emission (Em) and excitation (Ex) spectra and the linear relationship of optoacoustic signal and absorbance for **(a, b)** sfGFP, **(c, d)** mKate2, **(e, f)** mCherry, **(g, h)** HcRed, **(i, j)**tdTomato, **(k, l)** smURFP and (**m, n**) IRFP720 (IRFP). Representation similar to Figure 2.

Several explanations are possible for the higher PGE of the blue shoulder observed in most of the proteins. i) The blue shift could point to an involvement of the neutral state of the chromophore (commonly called an A state, as opposed to the anionic B form). For several FPs, fast (ps) ESPT from the excited state of the neutral form (A*) to an excited intermediate state (I*) are possible (Cotlet et al., 2002; Miyawaki, 2004). It is possible that the deexcitation from this state results in vibronic relaxation and a higher PGE. However, the excitation spectra for sfGFP, mKate, and mCherry have shapes that are similar to the optoacoustic spectra, all with significant pronunciation in the shorter wavelength range. Additionally, the fluorescent lifetimes for sfGFP exciting at 420 and 490 nm show only a slightly longer lifetime at 490 nm (2.77 vs. 2.37 ns), suggesting similar deexcitation compared to the B form. ii) For HcRed, our data is in agreement with the theory of a mixture of cis-trans isomers, with a non-fluorescent, neutral trans-isomer centered at around 570 nm and a slightly fluorescent cis-isomer centered at 590 nm (Mudalige et al., 2010; Wilmann et al., 2005). The high PGE from the peak at 570 nm could arise from the non-fluorescent properties of the trans-isomer and, independently, the tetrameric character of HcRed promoting exciton coupling between nearby chromophores, as observed in other multimeric proteins(Visser et al., 2002). iii) Beyond our previous explanation, discrepancies between OA and absorption spectra have been observed in some proteins (mCherry, mNeptune and dsRed) before and explained by ground state depopulation (Laufer et al., 2013). In short, when a fluorescent molecule is excited by early arriving photons with the pulsed illumination used in OA (5 - 7 ns), and if the fluorescence lifetime of the molecule is long enough, later arriving photons have a lower number of ground state electrons for excitation. This effect primarily come into play for the strongly absorbing central transitions of an absorption spectra, leading to the “flattened” absorption peaks observed by Laufer and colleagues (Laufer et al., 2013). Since in our study the discrepancies are regularly found at the flanks of the spectra or only for certain peaks (e.g., aggregate peaks or protonated chromophores), we believe the phenomenon of ground state depopulation does not explain our results. Interestingly, however, we can see that excitation spectra in Figure 4 show ground state depopulation (loss of the spectral shape with a difficult to differentiate main peak). This might be possible since electrons that decay through the radiative pathway have slower rate constants than those that decay through internal conversion (ns vs. ps) (S. P. McGlynn, 1969). Thus, if ground state depopulation due to the intense short pulse irradiation is present, it will primarily affect the radiative transitions and thus the fluorescence excitation spectra.

Another peculiarity, which we observed by recording spectra under conditions regularly used for OA imaging, is that several proteins showed a level of photoconversion or photo-switching that was previously unknown. This was initially observed when comparing forward (420 to 900 nm) and reverse (900 to 420 nm) modes of collecting spectral data. For mCherry and tdTomato, the forward and reverse measurements produced different results, which suggests long-lived states induced by the illumination. The effects are different for the two proteins, but always show a long-lived change of the OA spectra. For mCherry, measuring from 420 to 900 nm results in a stronger blue shoulder than measuring in the opposite direction (**Figure 5a**). Surprisingly, the absorption spectrum of mCherry does not change accordingly (**Figure 5b**), resulting in an increase of the PGE of the shoulder from 0.35 to 0.45. mCherry does not show evidence of phototransformation after continued illumination with 550 or 580 nm light (**Figure 5b**). For tdTomato, in contrast to mCherry, the shoulder increases when imaged from 900 to 420 nm with a change in PGE for both peaks (**Figure 5d and 4j**).

**Figure 5.**
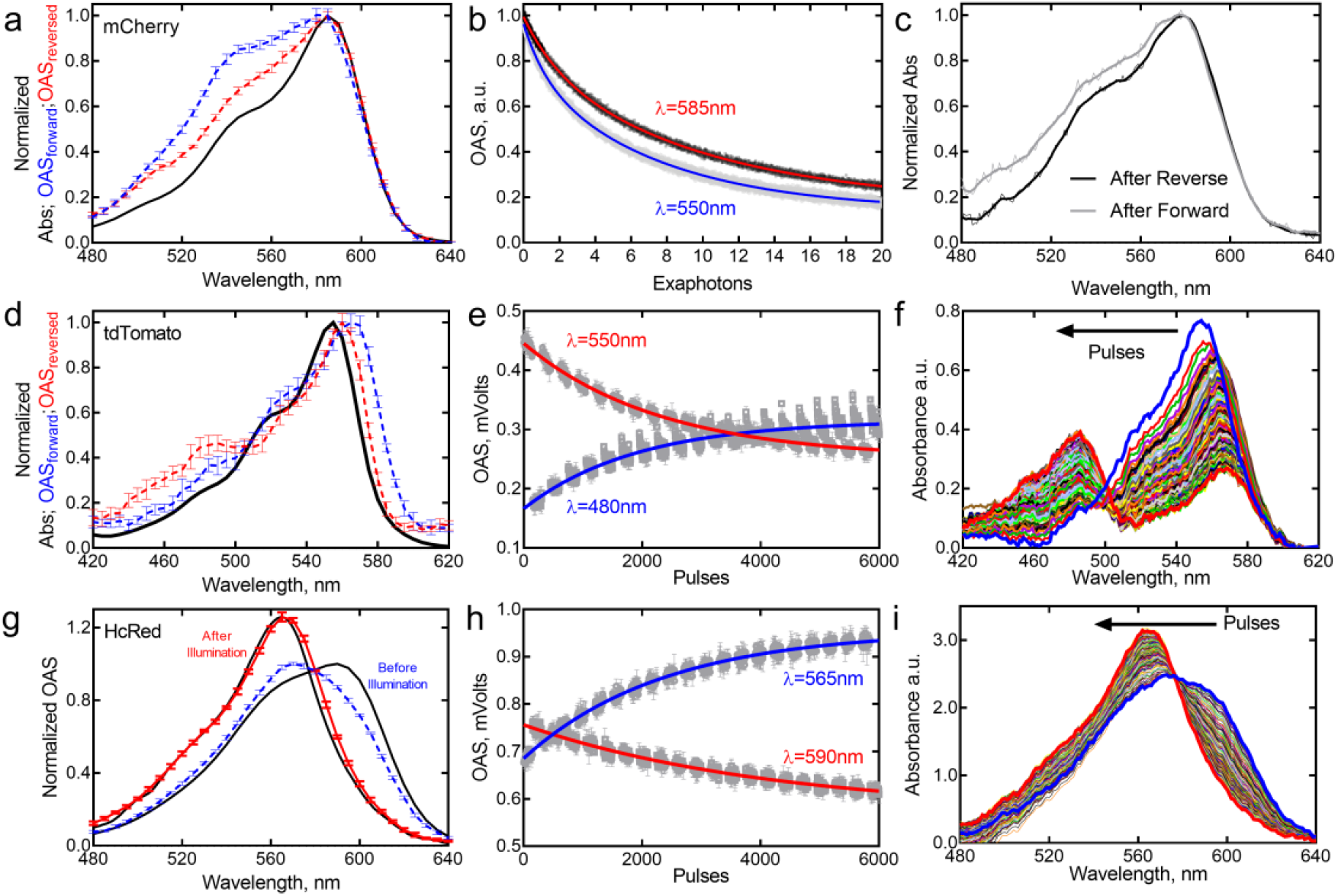
Photoswitching, -conversion or bleaching effects in mCherry, tdTomato and HcRed. Shown are the OA spectra measured from 900 to 420 nm (reverse, red) or 420 to 900 nm (forward, blue) along with the absorption spectra in the non-illuminated state (a, d and g). mCherry absorption spectra before and after OA spectral measurements (c) and after multiple pulses of 550/480nm and 590/565nm light for tdTomato and HcRed, respectively (f and i). The temporal development of the OA signals at the two main peaks (b, e and h).

Surprisingly, the maxima of the OA spectra also shift. Alternating illumination between 480 nm and 550 nm shows how the absorption spectra changed with an increase of the band at 480 nm, which can be attributed to the neutral chromophore in those proteins (**Figure 5e and 5f**). Both mCherry, as well as tdTomato, are related to the photo-switchable proteins rsCherry and rsCherryRev (Stiel et al., 2008), which primarily show fluorescence based photo-switching, but also some level of photochromism (more so for rsCherry). It could be that the effects in mCherry and tdTomato under strong pulsed laser illumination are similar to photo-switching events, although we see no dark-relaxation, typical of reversible photo-switching.

For the far-red protein, HcRed, continuous illumination at 565 nm did not produce a decrease in intensity (bleaching) as expected, but an increase in signal – an effect also observed for rsCherry (Stiel et al., 2008). However, in contrast to rsCherry, we did not observe an inflection point, even after 60,000 pulses. Alternating illumination between 565 nm and 590 nm results in a reduction in the signal at the fluorescence peak at 590, but an increase at the optoacoustic peak at 565nm; the absorption spectra show the same kinetics (**Figure 5g-i**). Trans to cis-isomerization of HcRed has been previously induced by heating (Mudalige et al., 2010), but until now, there was no report of light-induced changes in this protein. A published crystal structure of HcRed suggests inherent chromophore flexibility with the trans nonplanar isomer responsible for the shoulder at 570 nm, and cis-isomer responsible for the peak at 590 nm, with both conformer in equilibrium(Wilmann et al., 2005). In order to validate our proposed mechanism, we decided to crystalize our photoconverted version of HcRed. Our structure shows that the inherent chromophore flexibility is not present anymore after pulsed illumination (**supplementary Figure 7**). The new structure only has a trans-nonplanar chromophore that is tilted and stabilizes by new H-bond with a twisted His174. Our structure corroborates that trans-isomer is responsible for the optoacoustic peak at 565 nm. While such phenomena are interesting from a perspective of FP photophysics, they can also be useful for OA imaging, since the transitions can be used similarly as photo-switching for locked-in OA detection (Mishra et al., 2019). In this regard, tdTomato is of especially high interest, since a vast number of established lines and transgene animals exist with this protein due to its excellent properties for fluorescence imaging. It is all-the-more astonishing because in our measurements, tdTomato showed a high PGE of 0.8, which does not agree with its long fluorescence lifetime of around 3.6 ns or its reported quantum yield of 0.66. This discrepancy remains to be elucidated, but is favorable in this particular case for both OA and fluorescence imaging.

Biliverdin binding proteins are of critical importance in optoacoustic imaging, as their absorption in the near-infrared region gives them superior penetration depth and spectral separation for endogenous contrast. Until now, only two classes of near-infrared biliverdin binding proteins have been reported: bacteriophytochromes and phycobiliproteins (Rodriguez et al., 2016; Shcherbakova and Verkhusha, 2013). IRFP720 and smURFP are examples of both classes. In both cases, the optoacoustic spectra match the absorption spectra, with PGEs of 0.56 and 0.86 for smURFP and IRFP720 at their primary OA peaks, respectively (**Fig. 4k and m**). The lower PGE of smURFP correlates with its higher quantum yield 0.2, versus 0.06 for IRFP720. Additional reductions in PGE could come from faster bleaching and enhanced transient isomerization, a phenomenon already reported for biliverdin (Braslavsky et al., 1983a, 1983b). The excitation spectrum of smURFP shows evidence of ground state depopulation; even more so than for the classical proteins, ground state depopulation is not apparent in the optoacoustic spectrum (**Figure 4k**). Instead, IRFP720 shows a shoulder whose transition is less excitable and has a higher PGE (0.97, **Figure 4n**).

## CONCLUSION

Our first results using the MLS showed a wealth of insights into the behavior of a number of well-used dyes and proteins. Describing the exact changes in the photophysics of OA signal generation upon the aggregation of MB, Rh800, and ICG will afford new applications for sensors that exploit their environment-dependent aggregation. For example, ratiometric analysis of dye aggregation under different physiological conditions. A similar strategy is used in measurements of blood oxygenation with two wavelengths (Choi et al., 2016).

Furthermore, determining which transitions are most effective for OA imaging enables dyes like Alexa Fluor 750 to be employed more effectively for this modality. Finally, our results on ICG shed light on the complex behavior of this molecule, which is so regularly used in OA studies (Beziere et al., 2015; Wilson et al., 2018). Cognizance of the strong dependence of PGE on the aggregation of dyes could help researchers to analyze concentration-dependent measurements of ICG more accurately. On the protein side, the switching or conversion behavior of mCherry, tdTomato, and HcRed is extremely interesting for GFP-like protein research and warrants further study; however, it also has an immediate impact on employing those proteins in OA applications. In particular, tdTomato – for which there are numerous animal models from fluorescence studies – can be used more effectively for OA measurements by exploiting the transition from 550 nm to 480 nm and using a dual-wavelength ratio analysis to identify the protein-label in tissue with a strongly absorbing background.

Lastly, we could show that accurate measurements for OA labels are a prerequisite for their understanding, their future design, and especially their comparison. Regarding this, the use of standards like BBN for novel OA contrast agents to come is highly important for comparability, and for empowering researchers to make strategic choices on what labels to use in their experiments.

## MATERIAL AND METHODS

### Preparation of dyes

Dyes were dissolved at 5 mg/ml in DMSO and frozen; previous to measurement, a stock solution of the dyes 1% v/v was prepared in 10% FBS. The dye was further diluted to provide different concentrations. The absorbance spectra of each concentration were measured several times during one hour to assure no change in the absorption spectra over time.

### Preparation of proteins

Proteins were expressed in Escherichia coli BL21 and purified by Ni-NTA affinity chromatography, followed by gel filtration on a HiLoad 26/600 Superdex 75pg column (Amersham Biosciences) in PBS buffer. Purified proteins were frozen immediately in liquid nitrogen and stored at −80 °C. Thawed proteins were centrifuged at 14,000 rpm for 45 min at 4 °C, and the supernatant was used for the measurements.

### Multi-modal laser spectrometer setup

Building upon previous art (Vetschera et al., 2018), we developed a multi-modal laser spectrometer (MLS, Figure 1b) that allows us to record all three modalities (optoacoustics, absorbance, fluorescence) simultaneously in the wavelength range between 420-1000 nm. Figure 1b schematized the optical system consisting of: 1. A diode-pumped solid-state optical parametric oscillator (DPSS OPO, SpitLight DPSS250 OPO, Innolas Laser GmbH, Germany) laser covering wavelength range 420 - 2000 nm with a 10-50 Hz pulse repetition rate (PRR), a pulse width at full width at half maximum *τ_FWHM_* =7 ns, and a <1 nm linewidth of specific wavelength output. The OPO signal beam of 420-709 nm is output with horizontal polarization and idler beam 710-2000 nm with vertical polarization (switching controlled by a stepper motor and a microcontroller Arduino Uno via USB); 2. A pulse energy control to maintain constant pulse energy, which is measured using pyroelectric sensor head PE10-C and USB readout power meter Juno (Ophir-Spiricon LLC, USA), and controlled by using a combination of a motorized half-wave plate (HWP, AHWP05M-600; mounted on motorized rotation stage PRM1Z8; Thorlabs Inc., USA), a polarizing beam splitter (PBS052, Thorlabs Inc., USA), and a beam dump; 3. An optical filter wheel that switches between long-pass (LP, cut-off wavelength at 700 nm to allow for OPO idler beam) and short-pass (SP, cut-off at 800 nm for signal beam) edge filters that are switched by an Arduino Uno via a stepper motor, in order to remove the possible leakage of signal/idler at output beam; 4. An electrically controlled diaphragm shutter (SHB05T, Thorlabs Inc., USA) that is triggered by an Arduino Uno; And 5. a fiber bundle and its coupling lens (numerical aperture, NA 0.22; 430 fibers bundled in *ϕ*4.6mm; CeramOptec GmbH, Germany) to deliver light to a deionized water chamber which is determined for optimal coupling of optoacoustic wave.

In the water chamber, of which the walls were painted with black, we custom-built a holder to fix the relative position of all components, including: 1. The IBIDI chips (μ-Slide I Luer, 200 μm channel thickness, ibidi GmbH, Germany) for sample and reference materials (water solution of Indian ink). The in-line reference chip is inserted in the beam path to remove the influence of laser pulse energy fluctuation of the sample spectrum. 2. Multi-mode fibers and collimating lens that deliver broadband light from a deuterium tungsten lamp (DH-2000-BAL, Ocean Optics, USA) and collect the transmitted light through the sample to a Vis-NIR spectrometer (USB4000-VIS-NIR, Ocean Optics, USA). Both the illumination and the detection fiber were attached to a collimator lens rotated by 45° to the optoacoustic illumination-detection axis; And 3. an ultrasound transducer (V382-SU, central frequency 3.5 MHz, focus length 38.1 mm, diameter 13 mm) that simultaneously detects optoacoustic wave generated from reference and sample chips. The detected temporal optoacoustic signals were amplified by a 10-60 dB variable-gain amplifier (DHPVA, FEMTO Messtechnik GmbH, Germany) and sampled by a 12-bit digitizer (DAQ, Gage CSE1222, DynamicSignals LLC, USA). At each wavelength 3-10 sequences are acquired for signal averaging.

The data acquisitions using DAQ and Vis-NIR spectrometer were synchronized using either the trigger from the laser or the output of a photodetector (Si Biased Detector, DET36A/M, Thorlabs Inc., USA) induced by stray light in laser beam path. All data acquisition, processing and setup synchronization are controlled by our custom-built programs using Matlab 2015b (The MathWorks Inc., USA).

### Synchronization of signals measurement

Empowered by the MLS spectrometer, four spectra of a sample can be obtained simultaneously: optoacoustics, absorbance, fluorescence excitation, and emission. 200 μL samples were filled in IBDI chips.

Since fluorescence and optoacoustic signal was excited by the same laser pulses, both can be recorded simultaneously. We set the Vis-NIR spectrometer in external triggering mode for excitation/emission spectrum measurement and shared the trigger as optoacoustic recording. When measuring optoacoustic spectrum and fluorescence spectra, the electrical shutter is opened, while the shutter to the lamp of Vis-NIR spectrometer is closed, and the ultrasound transducer, DAQ, and spectrometer (configured to external triggering mode and fixed integration time of 10-90 ms) are triggered by the laser or a photodiode to acquire data. Due to the coincidence of sweeping excitation laser and fluorescence emission, there was overlapping region of the two in a direct recording. To extract the excitation spectrum only, for example, from the recordings, we used the normalized readings of three different emission wavelengths at each excitation wavelengths. When the excitation wavelength was overlapping with one of the selected emission wavelengths, we ignored the current emission reading and used the extrapolated value averaged from the other two emission wavelengths. Similar method was adopted to extract the emission spectrum from such overlapping recordings.

When measuring absorbance, only the Vis-NIR spectrometer acquires data (configured to circular-buffered acquisition mode using integration time that uses 90% of the full pixel well depth when making dark/no sample measurement), and laser light is blocked by the closed shutter and lamp shutter is opened to deliver broadband white light to the sample. The setup has a time delay for the absorption measurement of at least 110 ms after the laser pulses due to shutter responses.

### Lookup table for constant pulse energy

One challenge in measuring with high-fidelity on common dyes/proteins based chromophores is that they suffer from photobleaching and such processes are usually wavelength dependent. Constant pulse energy was ensured by the use of half-wave plate on motorized rotation stage and a polarizing beam splitter; using a lookup table and adapting the polarization with the halfwave plate, we kept the power constant at 1.2 mJ for the proteins measurements and 1 mJ for the dyes, with root-mean-squared error (RMSE) 2% over the whole illumination spectrum. We confirmed the linearity and stability of the optoacoustic signal generated by BBN and NiCl_2_ in the range of fluences used in our experiments.

Thus to ensure a controlled constant delivery of pulse energy to the sample across the 420-1000 nm range, the operation of MLS starts with a pulse energy lookup table measurement. Supplementary Figure 1 shows a complete flow chart on our high-precision laser pulse energy lookup table measurement method. The lookup table is intended to reflect the sinusoidal oscillation relation between the angle difference of HWP/PBS and the pulse energy of OPO at certain wavelengths. Therefore, we used a measure-and-fit scheme (Eq. 3) to fasten the speed of lookup table measurement.

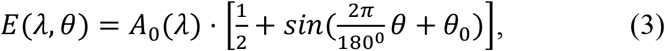

where *E*(*λ, θ*) is the pulse energy measured at laser wavelength when HWP is at angle *θ*; *A*_0_(*λ*) is the maximum pulse energy that is achievable only when HWP angle is at *θ*_0_ that coincides with the transmission polarization of PBS.

According to Eq. 3 (shown in Supplementary Figure 1), we first measured the energy at one OPO signal wavelength, e.g., at 420 nm with HWP angle rotating with step 2° across 72° angles (in order to include one peak and one trough). Then we fitted the energy-angle curve to the parametric sinusoidal model as indicated by Eq. 3, to find the phase angle *θ*_0_ where the peak energy appears. Fixing HWP at the peak phase angle *θ*_0_, we continued to measure the peak energy *A*_0_(*λ*) of the rest signal wavelengths, and based on which we extrapolated the sinusoids model. We showed in Supplementary Figure 2a an exemplary laser pulse energy lookup table measured using our measure-and-fit based scheme. The same procedure was applied to OPO idler wavelengths (≥ 710 nm) to obtain a crude estimate of the lookup table. But due to the fluctuation of OPO laser pulse, especially at wavelengths around 710 nm where signal/idler beam are separated, the extrapolated lookup table required still correction. Aiming at a pulse energy of e.g. 1.0 mJ at all wavelengths, we measured the pulse energy back using the angles suggested by the lookup table targeting at 1.0 mJ, and obtained the ratio of the targeted energy and the new measurements. Then we multiply this ratio by the amplitude *A*_0_(*λ*) to scale the lookup table proportionally. Usually, less than 5 iterations of this lookup table correction would enable a RMSE of <2% at a specified pulse energy, as shown in Supplementary Figure 2b.

### Principle of in-line correction

Another challenge resides with the pulse-to-pulse fluctuation at given wavelength after homogenizing the laser pulse energy, which can range from a few percent to more than 30% as at the transition wavelengths of signal/idler output (Supplementary Figure 2b). To minimize the influence of such fluctuation, we designed an in-line reference using water solution of Indian ink in optoacoustic sensing. The resulting OA signals from both reference and sample are collected by the ultrasound transducer in the same temporal sequence. As shown in Figure 1b, laser light fluence *φ*_0_ with fluctuation Δ*fl* traveling a water path *d_FR_* from fiber to reference chip pass through the low concentration of ink with a light fluence *φ_R_* according to Beer-Lambert’s law:

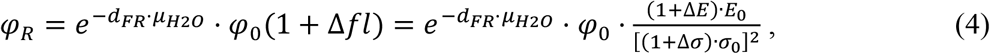

where *μ_H2O_* is water absorbance. It is important to note that the pulse-to-pulse light fluence fluctuation Δ*fl* manifests as both a peak power (*E*_0_) variation (Δ*E*) and a pulse width (*σ*_0_) variation (Δ*σ*), assuming Gaussian temporal profile. The pulse width variation (sub-nanosecond for 7 ns pulse width we used) is a term cannot be corrected using a slow-responding power meter or photodiode for optoacosutics, but only possible by using high-speed photodiode (<100 picosecond) and fast acquisition system (>20 GHz), which is a costly. Instead, we used optoacoustic reference. The transmitted light after reference chip with absorbance *μ_aR_* and path *d* same to channel thickness of IBIDI chip then travels a water path *d_RS_* from reference chip to sample chip. The incidence light fluence to the sample, *φ_S_*, is:

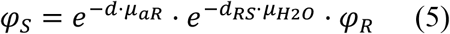

The photoacoustic signal generation induced by optical absorption and the subsequent thermal expansion can be described by a simplified model (Beard, 2011), *p_0_* = *Γμ_a_φ*, assuming a linear relation between absorption coefficient and light fluence in low absorbance and low scattering sample (Cox et al., 2012), *Γ* is the Grüneisen coefficient indicating the conversion efficiency from heat to pressure wave. Therefore, the optoacoustic wave generated by the reference chip is:

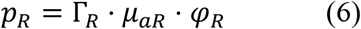

Similarly, the sample generated optoacoustic wave is:

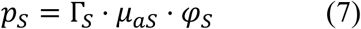

Dividing sample optoacoustic signal by reference can cancel the influence of laser pulse-to-pulse fluctuation if substituting in the light fluence to reference (Eq. 4) and sample (Eq. 5).

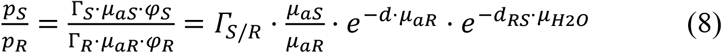

As a consequence, Eq. 8 indicates that fluctuation correction renders a mixed spectrum of the reference and sample. The first term on the right-hand side of Eq.8 is a ratio of Grüneisen coefficients of sample over reference. The second term is absorption coefficient ratio of sample over reference. The third term indicates the spectral coloring effect (Tzoumas et al., 2016) (light fluence alteration) by the in-line reference. The fourth term represents the spectral coloring by the water path between reference and sample chips.

There are ways to account for the effect of above-mentioned terms. The absorption coefficient *μ_aR_* of reference solution can be obtained by measuring its absorbance A (cm^−1^) and converting using *μ_aR_* = *ln*(10) · *A*, assuming low scattering by a low concentration (i.e. absorbance) of ink. Thus multiplying the spectrometer measured *μ_aR_* and Eq.8 would remove the influence by ink spectra. Furthermore, although the effect is minor, the spectral coloring effect by ink can be removed, given the thickness of IBIDI chip channel thickness. Similarly, given water path *d_RS_* between reference and sample chip and water absorption coefficient (data adapted from http://www.spectra.arizona.edu/), the spectral coloring effect by water can be removed. The corrected optoacoustic signal at wavelength as a result is:

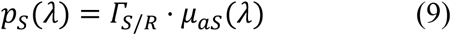

where the Grüneisen coefficient ratio *Γ_S/R_* is not supposed to be wavelength dependent thus a scaling factor of optoacoustic signal amplitude. For reproducible comparison across various samples, if we use ink as reference for all the samples, the Grüneisen coefficient ratio still reflects the heat conversion efficiency of samples.

### Data process for multi-modal measurement

Supplementary Figure 3 shows the program for data acquisition and data process in multimodal laser spectroscopy. According to our in-line referencing to correct for the laser pulse-to-pulse fluctuation, we applied similar process to fluorescence emission data as to optoacoustic data.

The computation of the spectra (blue shading part showed in Supplementary Figure 3) is done as described in the following sequential steps: taking a sample of e.g. NiCl_2_ with 0.5 OD at 720 nm. The results are shown in Figure 1c, providing an examination on the improvement brought by each of the processes in comparison with the raw spectrum - Raw (Max) in Figure 1c - which is acquired by taking simply the maximum value measured at each wavelength upon homogenized laser pulse energy. Matching the absorbance spectrum, the Raw spectra showed a coefficient of determination R_2_ of 0.988 and a standard deviation 7.9% between repeated recording, owing to the 2%-RMSE laser energy control (Supplementary Figure 2).

1. *‘Filtered’*. We firstly filtered the acquired temporal sequence by using a fourth order digital bandpass finite impulse response filter in Fourier frequency band 0.1-10 MHz respecting the bandwidth of our transducer, a 1-D digital wavelet transform (decomposition level=5) based denoising method to reduce thermal noise level, a 3rd order Savitzky-Golay filter with smoothing length corresponding to 20 MHz in our DAQ (200 MSamples/second) to reduce digitization error.
2. *‘Hilbert’*. Hilbert transform is used to detect the peak envelope of temporal sequence. The optoacoustic spectrum relies on the maximum signal in the sequence at the position (time-of-flight) of sample and reference. We detected the peak signals by summing up all sequences measured at all wavelengths, then finding the locations of the peaks of sample and reference respectively, and using the peak locations as center point to take averaged values of adjacent 3-5 sample points from sequence of each wavelength. A better signal-to-noise ratio (SNR) of optoacoustic spectra for low absorbance sample was achieved using this way than taking area under the curve of signal envelope.
3. *‘-Dark’*. We subtracted from the raw signal with the root mean square of a dark background signal that was measured with no laser irradiation before each spectral measurement to reduce the noise floor and thus improve the detection sensitivity of low concentration sample.
4. *‘Pulse Fl.’* To reduce the influence of laser pulse-to-pulse energy fluctuation, we divided the sample spectrum acquired at each laser pulse by the corresponding reference spectrum. But this step introduces spectra of ink into the sample spectra.
5. *‘-Ink Spect.’* To remove the ink spectra, we multiplied the fluctuation corrected spectra with normalized spectra of ink to recover the sample spectra.
6. *‘-Coloring’*. Notwithstanding that it is minor effect in 420-900 nm range induced by the spectral coloring by ink (short absorption path) and water path between reference and sample (water absorbance at 900 nm is 0.025 cm-1), we corrected for spectral colorings. The improvement was neither reflected in the coefficient of determination R_2_ nor standard deviation of measurements, but was instead shown as a higher fidelity in the peak value at 420 nm as shown in Figure 1c.

## ASSOCIATED CONTENT

### Supporting Information

The Supporting Information is available free of charge

### Author Contributions

J.P.F.W. and Y.H. build the MLS with help from P.V. in early stages. Y.H. programmed the measurement routine. J.P.F.W., Y.H. and K.M. conducted measurements. R.J. solved the structure. A.C., D.N. and V.N. contributed to the manuscript. A.C.S. J.P.F.W. and Y.H. wrote the manuscript and devised the project. All authors approved the manuscript. J.P.F.W. and Y.H. contributed equally.

### Funding Sources

Y.H. acknowledges the support from the China Scholarship Council via fellowship (CSC 201306960006). K.M. and A.C.S. acknowledge funding through Deutsche Forschungsgemeinschaft (STI 656/1-1).

## ACKNOWLEDGMENT

The authors want to address special thanks to Hong Yang, Dr. Ara Ghazaryan, Dr. Yu-Shan Huang, Dr. Christian Zakian, and to Dr. Jan G. Laufer for inspiring discussions.

