## Supplementary material for "Challenging a preconception: Optoacoustic spectrum differs from the absorption spectrum of proteins and dyes for molecular imaging": Supplmental Info

### Supplemental Information.

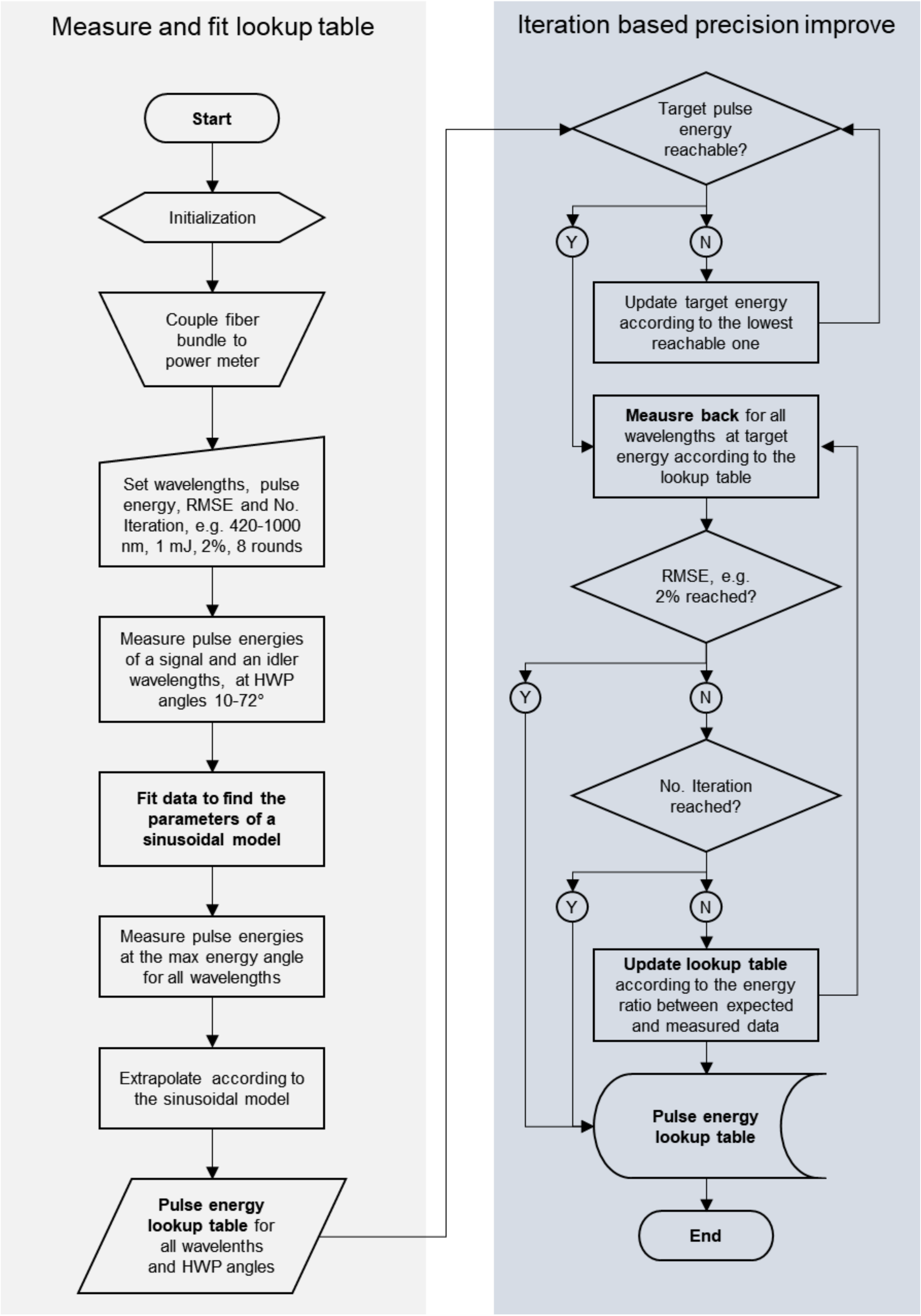


**Supplementary Figure 1** Flow chart of laser pulse energy lookup table measurement. In grey shading **(left)** is the measure-and-fit scheme measuring laser pulse energy controlled by rotary half-wave plate and polarizing beam splitter. In blue shading **(right)** is the iteration-based pulse energy precision improvement method.

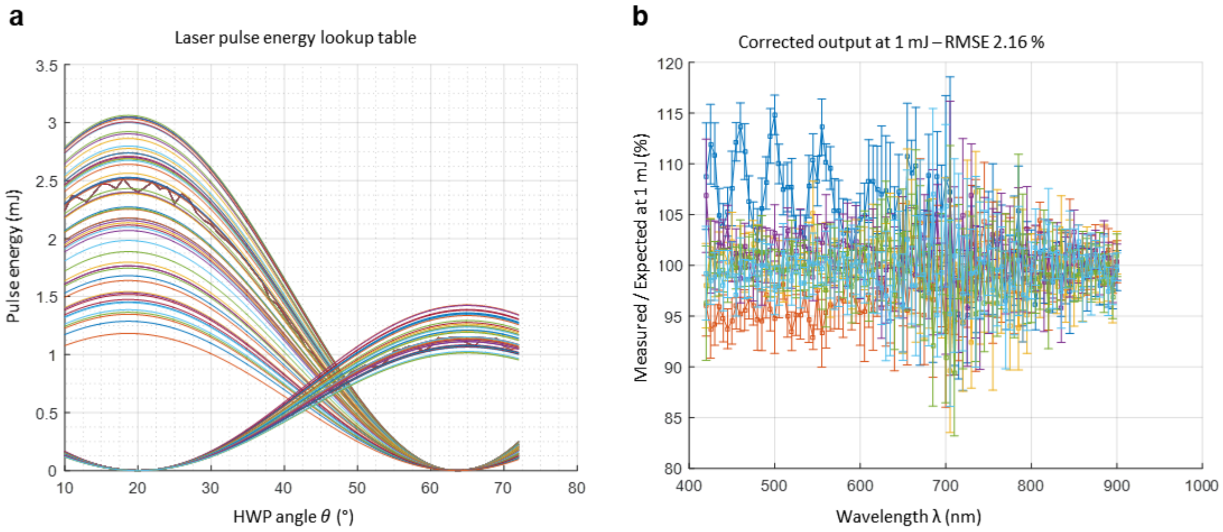


**Supplementary Figure 2 (a)** Measure-and-fit scheme measuring laser pulse energy controlled by rotary half-wave plate and polarizing beam splitter. **(b)** Iterated pulse energy lookup table correction.

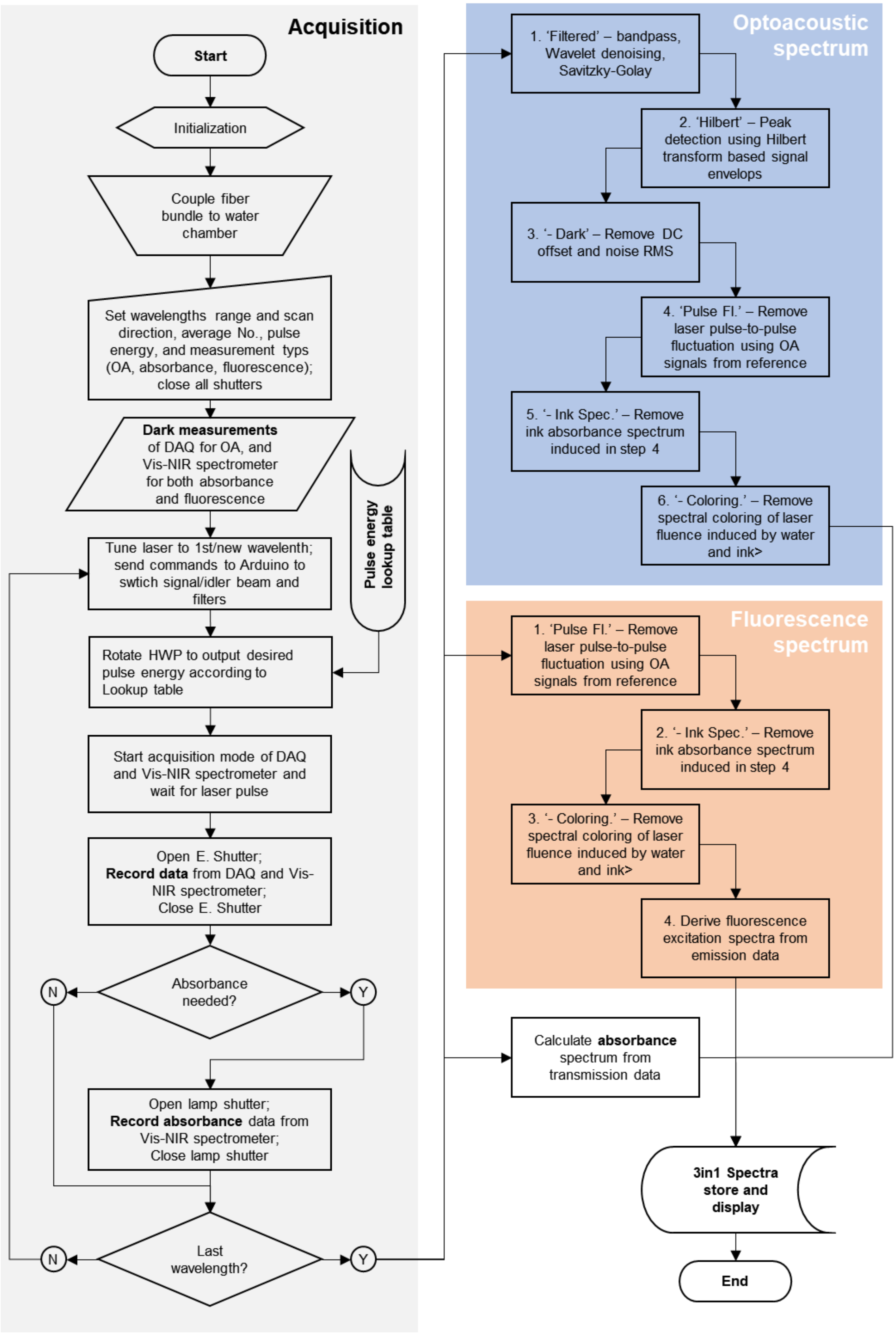


**Supplementary Figure 3** Flow chart showing the data acquisition (in **grey** shading, left) and data process scheme (right; in **blue** for optoacoustic spectrum; in **orange** for fluorescence excitation and emission spectrum; absorbance is showed without special marks). Data input showed as 'Pulser energy lookup table' is imported as a result of measurements showed in Supplementary Figure 1.

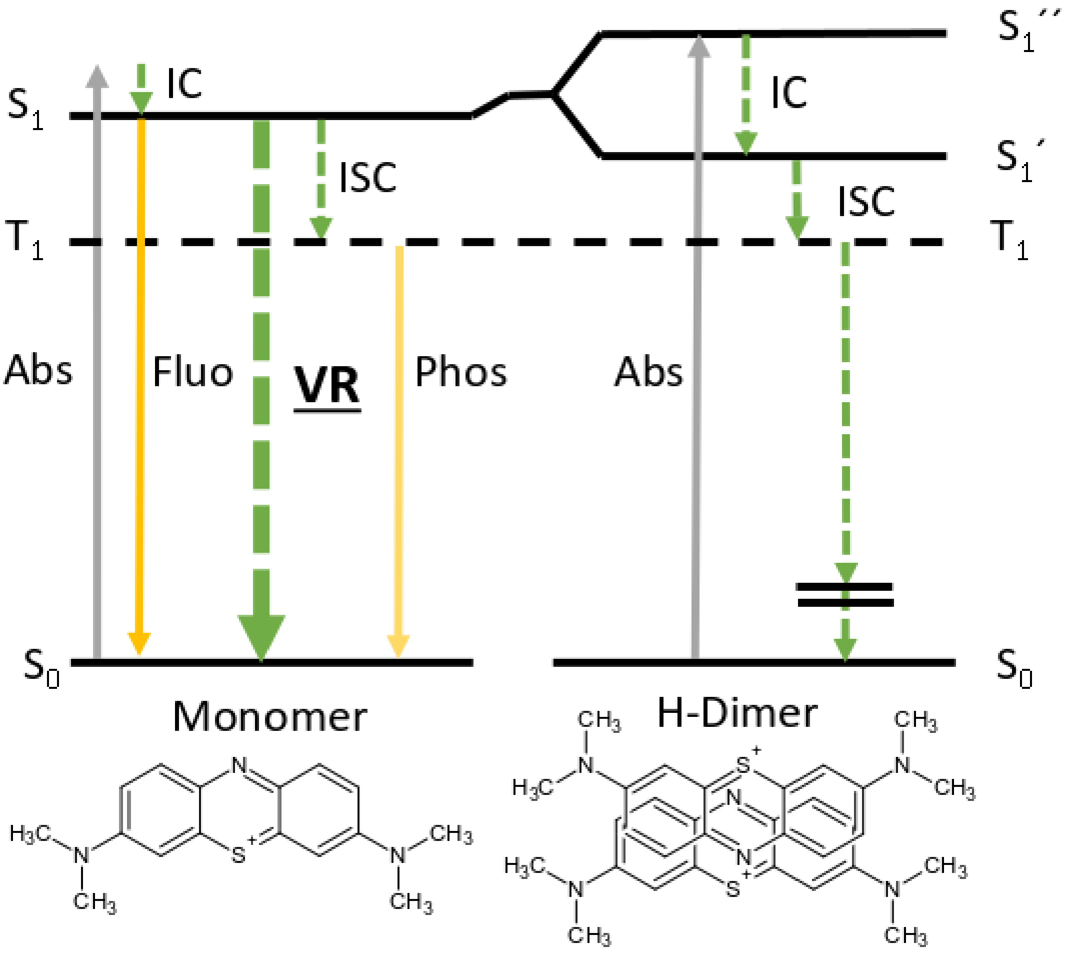


**Supplementary Figure 4** Jablonski energy diagram for monomers and geometrically perfect H-type dimers of Methylene blue. Plain lines represent radiative transitions, dashed lines represent nonradiative processes. Abs: absorbance $S_0$: ground state, $S_1$: singlet excited state, $T_1$: primary triplet excited state, $S'_1$ and $S''_1$: split exciton states, VR: vibrational relaxation, ISC: intersystem crossing, IC: internal conversion, Pho: phosphorescence, Fluo: Fluoresence. The double bar in the relaxation from $T_1$ indicates a chance for discontinuity in the transition diagram: the dimer triplet can undergo electron transfer. **Bottom**: possible configurations of MB monomer and H-type dimer. Modify from Kasha *et al.* 1965 [30] and Morgounova *et al.* 2013 [26].

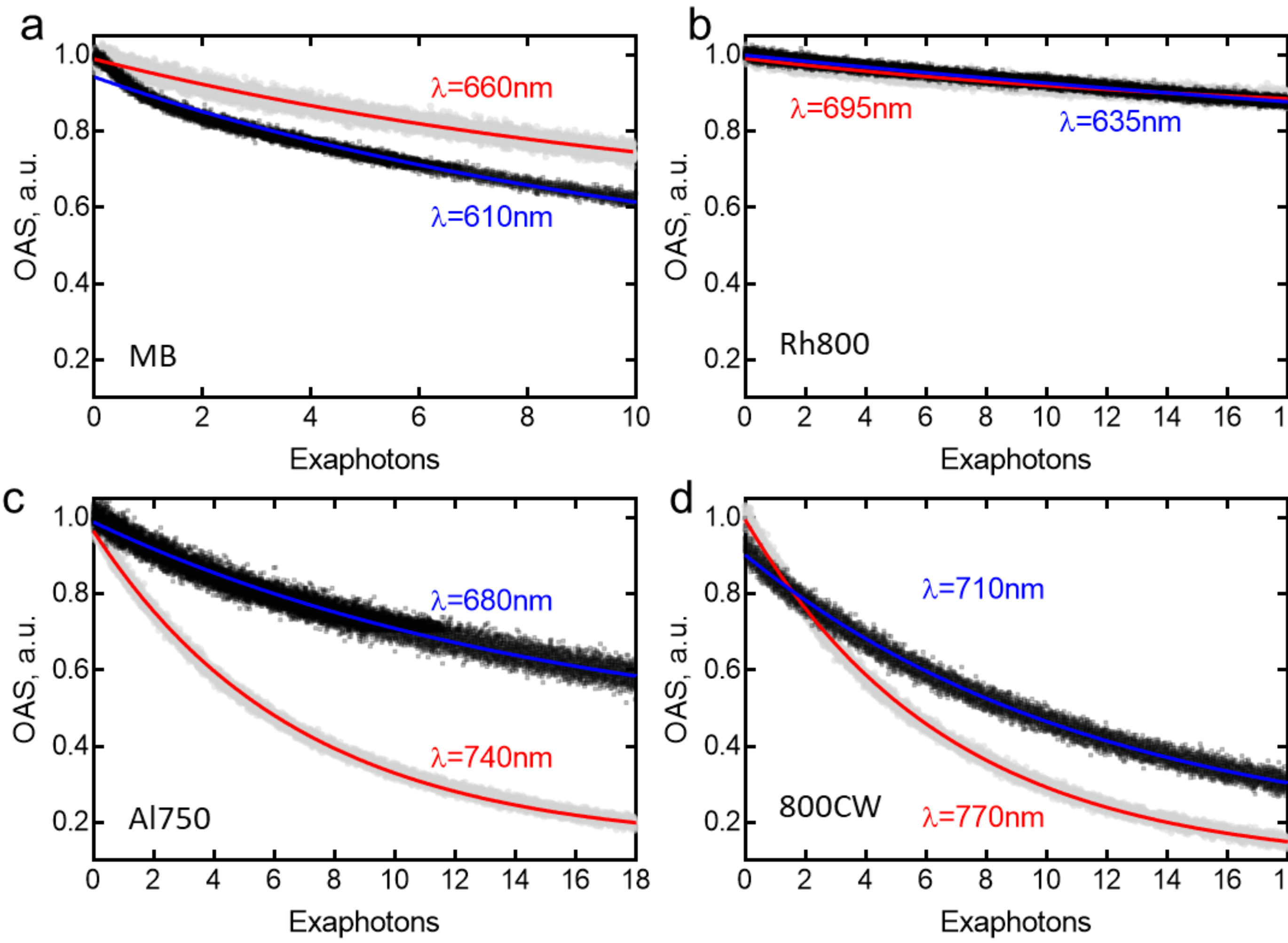


**Supplementary Figure 5** Bleaching measured as optoacoustic signal of (**a**) Methylene Bue - MB, (**b**) Rhodamine 800 – Rh800, (**c**) Alexa Fluor® 750 – Al750, and (**d**) IRDye 800CW at shoulder and main peak wavelengths.

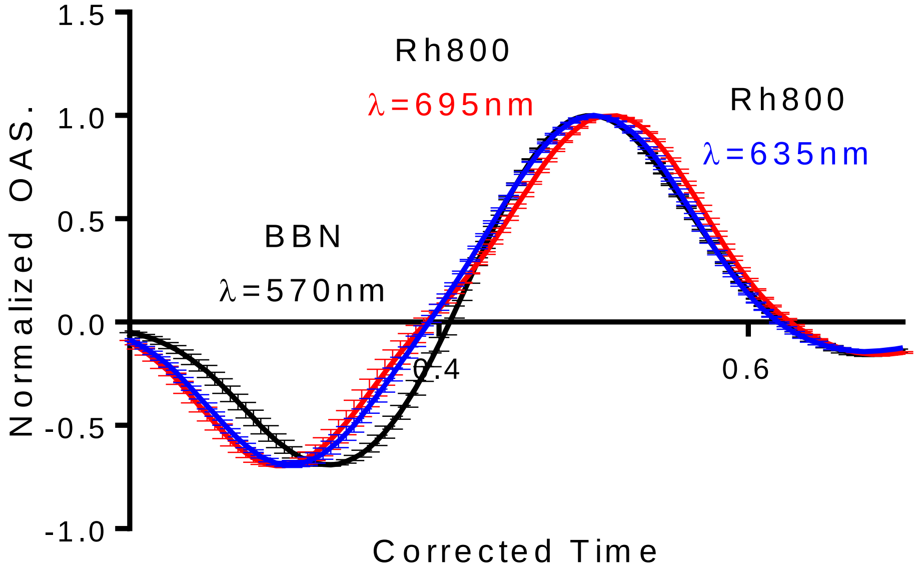


**Supplementary Figure 6** Optoacoustic signal for Rh800 at 695 nm and 635 nm; and BBN at 570 nm.

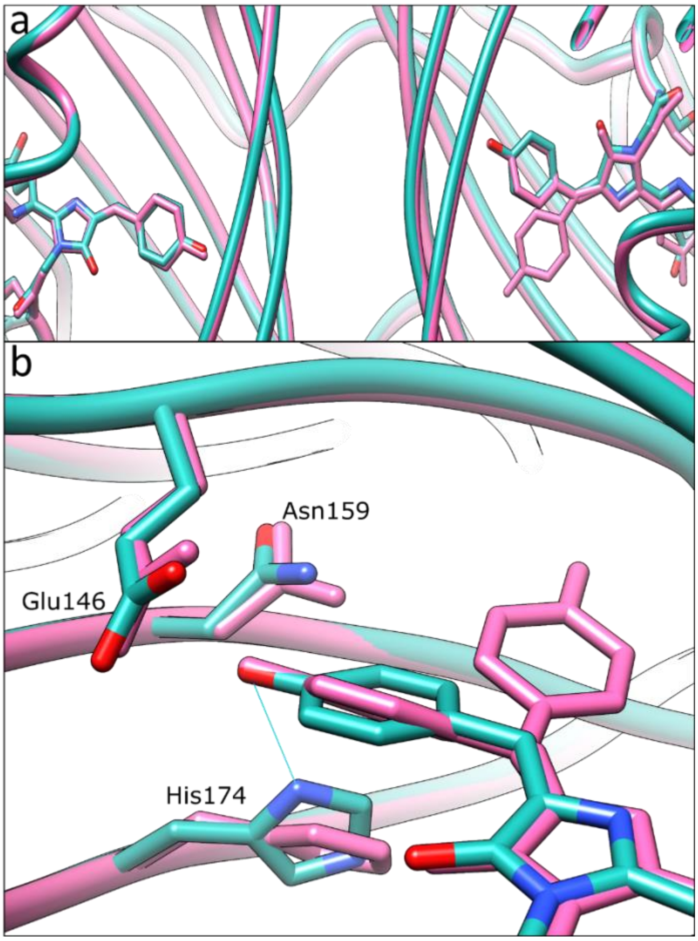


**Supplementary Figure 7** In blue structure of HcRed after 60000 pulses of 565 nm light and in pink structure of native HcRed (PDB code 1YZW): (**a**) dimer interface, and (**b**) Hydrogen bond between His174 and trans-nonplanar conformer in illuminated HcRed (blue).

**Supplementary Table 1** Characteristics numbers of dyes **(a)** and proteins **(b)** studied in this work.

#### Supplementary Table 1a

| | Absorbance Max1, nm | Absorbance Max2, nm | Optoacoustic Max1, nm | Optoacoustic Max2, nm | Excitation Max, nm | Emission Max, nm | Linear Fit at Max1 (**PGE**)[a] | Linear Fit at Max2 (**PGE**) [a] | Bleaching at Max1[b] | Bleaching at Max2[b] |
|---|---|---|---|---|---|---|---|---|---|---|
| MB | 615 | 665 | 610 | 665 | 665 | 680 | **0,61***X - 0,049 | **0,37***X - 0,073 | 0.73±0.01 | 0.85±0.01 |
| Rh800 | 630 | 690 | 635 | 695 | 690 | 705 | **1,14***X - 0,43 | **1,08***X - 0,40 | 0.95±0.02 | 0.96±0.02 |
| Al750 | 698 | 750 | 696 | 740 | 745 | 770 | **0,85***X - 0,04 | **0,72***X - 0,14 | 0.85±0.02 | 0.53±0.02 |
| 800CW | 710 | 775 | 710 | 770 | 770 | 789 | **0,86***X - 0,04 | **0,63***X - 0,02 | 0.63±0.02 | 0.5±0.01 |

#### Supplementary Table 1b

| | Absorbance Max1, nm | Absorbance Max2, nm | Optoacoustic Max1, nm | Optoacoustic Max2, nm | Excitation Max, nm | Emission Max, nm | Fluorescence Quantum Yield | Linear Fit at Max1 (**PGE**)[a] | Linear Fit AtMax2 (**PGE**)[a] |
|---|---|---|---|---|---|---|---|---|---|
| sfGFP | - | 490 | 425 | 490 | *** | 503 | 0.65 | **0,78***X - 0,01 | **0,34***X - 0,05 |
| mKate2 | 555 | 585 | 550 | 580 | 590 | 615 | 0.4 | **0,64**X+0.0 1 | **0,51**X+-0.03 |
| mCherry | 545 | 585 | 545 | 580 | 585 | 610 | 0.22 | **0.54***X + 0.01 | **0.36***X - 0.01 |
| tdTomato | 485 | 555 | 485 | 565 | 555 | 580 | 0.69 | **0.80***X - 0.02 | **0.83***X - 0.08 |
| HcRed | 570 | 590 | 570 | 590 | 590 | 635 | 0.015 | **0.97***X - 0.01 | **0.80***X - 0.03 |
| smURFP | 600 | 645 | 600 | 645 | 645 | 666 | 0.18 | **0.58***X - 0.01 | **0.55***X - 0.01 |
| IRFP720 | 650 | 705 | 650 | 700 | 705 | 720 | 0.06 | **0.97***X-0.09 | **0.86***X - 0.01 |

a. Linear Fit of absorbance versus corrected optoacoustic signal, PGE: Optoacoustic generation efficiency.

b. Bleaching rate after 5000 exaphotons.
